# ON/OFF domains shape receptive fields in mouse visual cortex

**DOI:** 10.1101/2021.09.09.459679

**Authors:** Elaine Tring, Konnie Duan, Dario L. Ringach

**Affiliations:** Department of Neurobiology, David Geffen School of Medicine, UCLA, Los Angeles, CA 90095; Harvard-Westlake School, Studio City, CA 91604; Department of Psychology, UCLA, Los Angeles, CA 90095

## Abstract

In higher mammals, thalamic afferents to primary visual cortex (area V1) segregate according to their responses to increases (ON) or decreases (OFF) in luminance^1–4^. This organization induces columnar, ON/OFF domains postulated to provide a scaffold for the emergence of orientation tuning^2,5–15^. To further test this idea, we asked whether ON/OFF domains exist in mouse V1 – a species containing orientation tuned, simple cells, like those found in other mammals^16–19^. Here we show that mouse V1 is indeed parceled into ON/OFF domains. Revealingly, fluctuations in the relative density ON/OFF neurons on the cortical surface mirror fluctuations in the relative density of ON/OFF receptive field centers on the visual field. In each cortical volume examined, the average of simple-cell receptive fields correlates with the difference between the average of ON and OFF receptive fields^7^. Moreover, the local diversity of simple-cell receptive fields is explained by a model in which neurons linearly combine a small number of ON and OFF signals available in their cortical neighborhoods^15,20^. Altogether, these findings indicate that ON/OFF domains originate in fluctuations of the spatial density of ON/OFF inputs on the visual field which, in turn, shapes the structure of receptive fields^10–13,21–23^.

## Main

We used two-photon imaging in mouse V1 to measure the responses of neurons in a cortical volume to sparse-noise stimulation (**Fig. 1a, Methods**). Each volume was sampled using 5 − 9 optical sections spaced 30 *μ*m apart (**Fig. 1b**). A standard data analysis pipeline comprised of image registration, cell segmentation, signal extraction and deconvolution steps, yielded the estimated spiking of neurons (**Fig 1c**). The centroid of the regions of interest (ROIs), along with the depth of the optical section, allowed us to assign each neuron a coordinate in cortical space, (*x*_1_, *x*_2_, *x*_3_) (**Fig 1b**).

**Figure 1.**
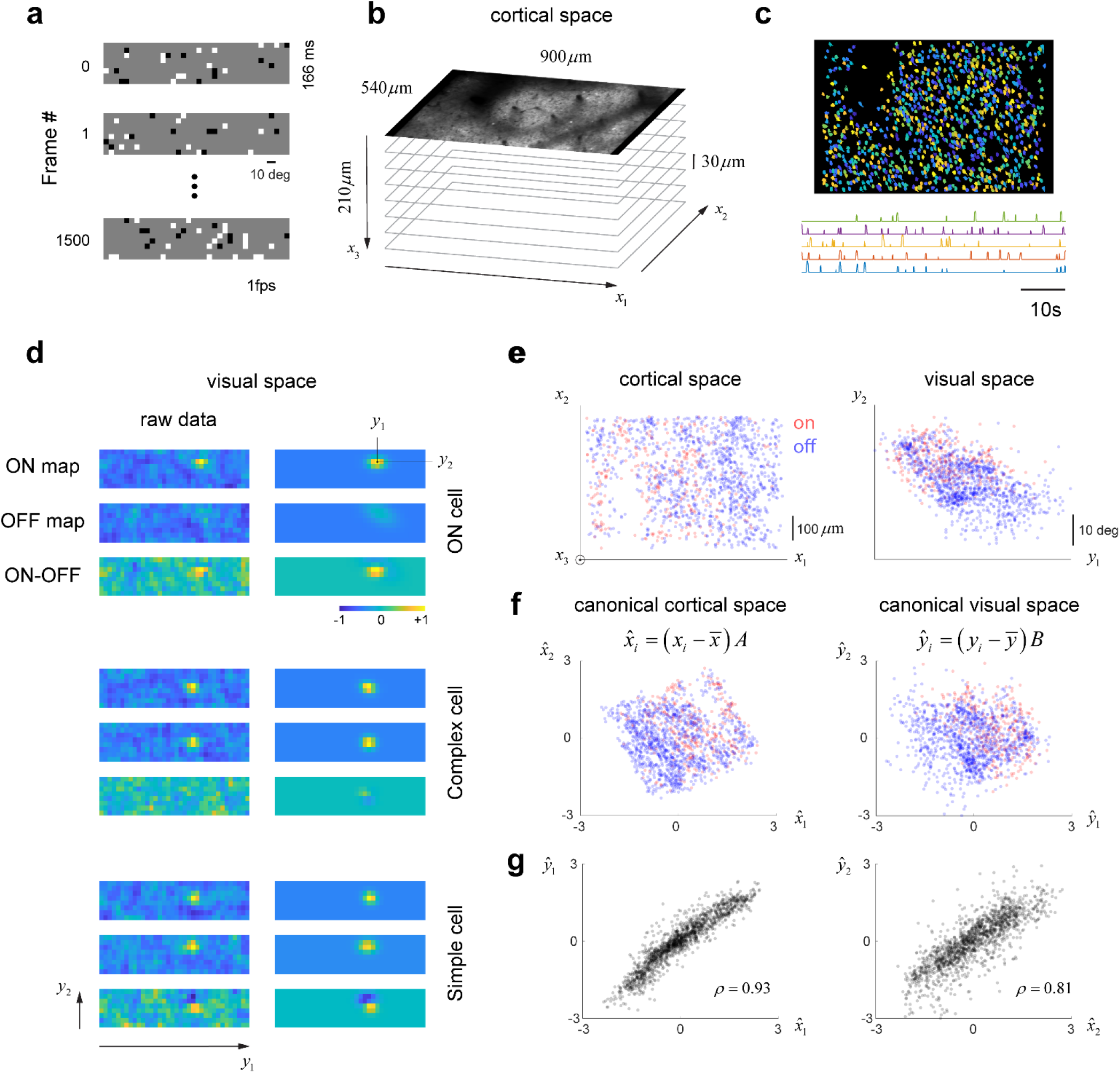
Representation of ON/OFF responses in cortical and visual space. **a**, Sparse noise stimulus. Images were flashed for 166ms and presented at a rate of 1 per second on a wide field screen. **b**, Volumetric sampling in primary visual cortex. **c**, Segmentation of regions of interest and sample traces showing spike inference from calcium signals. **d**, Raw receptive field measurements (left panels) and their Gaussian fits (right panels). **Top**, Cell with only a significant ON map. **Middle**, a complex cell with largely overlapping ON and OFF maps. **Bottom**, a simple cell with spatially displaced ON and OFF maps. In each case, ON and OFF maps are normalized jointly to the maximum absolute value attained by either of them. **e**, Distribution of ON (red) and OFF (blue) neurons in the native cortical and visual spaces. **f**, Same distribution now represented in canonical cortical and visual spaces. **g**, Correlation between the first pair of canonical variables 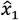 and 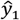 (left) and the second pair 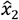 and 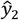 (right).

We computed the ON and OFF maps of each cell in the population by reverse-correlation (**Fig. 1d**, left panels, **Methods**). Cells that had only a statistically significant ON map with a single subregion were defined as ON cells; a similar definition applied to OFF cells. Cells with statistically significant ON and OFF maps were classified as simple or complex depending on the degree of spatial overlap between the maps (**Fig. 1d**)^24,25^. We fit two-dimensional Gaussians to ON and OFF maps to estimate their center locations (*y*_1_, *y*_2_) on the visual field (**Fig. 1d**, right panels). In simple cells, where the peaks of the ON and OFF maps are displaced in space, the difference between the ON and the OFF maps showed flanking subregions of opposite signs characteristic of their receptive field structure across mammalian species (**Fig. 1d**, bottom panels).

We analyzed the spatial distribution of ON and OFF cells on cortical and visual space (**Fig. 1e**). It was convenient, for reasons that will become clear soon, to bring these two coordinate systems into alignment using canonical correlation analysis^26^, which produced linear transformations of (*x*_1_, *x*_2_, *x*_3_) and (*y*_1_, *y*_2_) that were maximally correlated. The inclusion of cortical depth (*x*_3_) allowed us to compensate for slight departures of the objective from the surface normal. We represented each neuron in the population by its canonical coordinates in cortical space 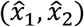 and its canonical coordinates in visual space 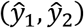 (**Fig. 1f**). In this example, the correlation between the first pair of canonical variables, 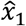 and 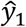, was *ρ* = 0.92, while the correlation between the second pair, 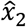 and 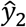, was *ρ* = 0.76 (**Fig. 1g**). These transformations ensure that the co-variances between 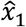 and 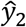, and 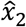 and 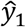, are zero (they are uncorrelated), and the variance of each canonical variable equals one.

### Mouse V1 is parceled into ON/OFF domains

We used kernel density techniques to estimate the probability distribution of ON and OFF cells in canonical cortical space, denoted by 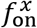 and 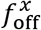 respectively (**Methods**). Similarly, we estimated the probability distribution of ON and OFF receptive fields centers in canonical visual space, yielding 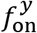 and 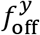. To detect fluctuations in the spatial distribution of ON and OFF cells in cortical space, we calculated the difference 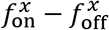. Similarly, to detect fluctuations in the distribution of ON and OFF receptive field centers in the visual field, we calculated the difference 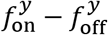 (**Fig. 2a**).

**Figure 2.**
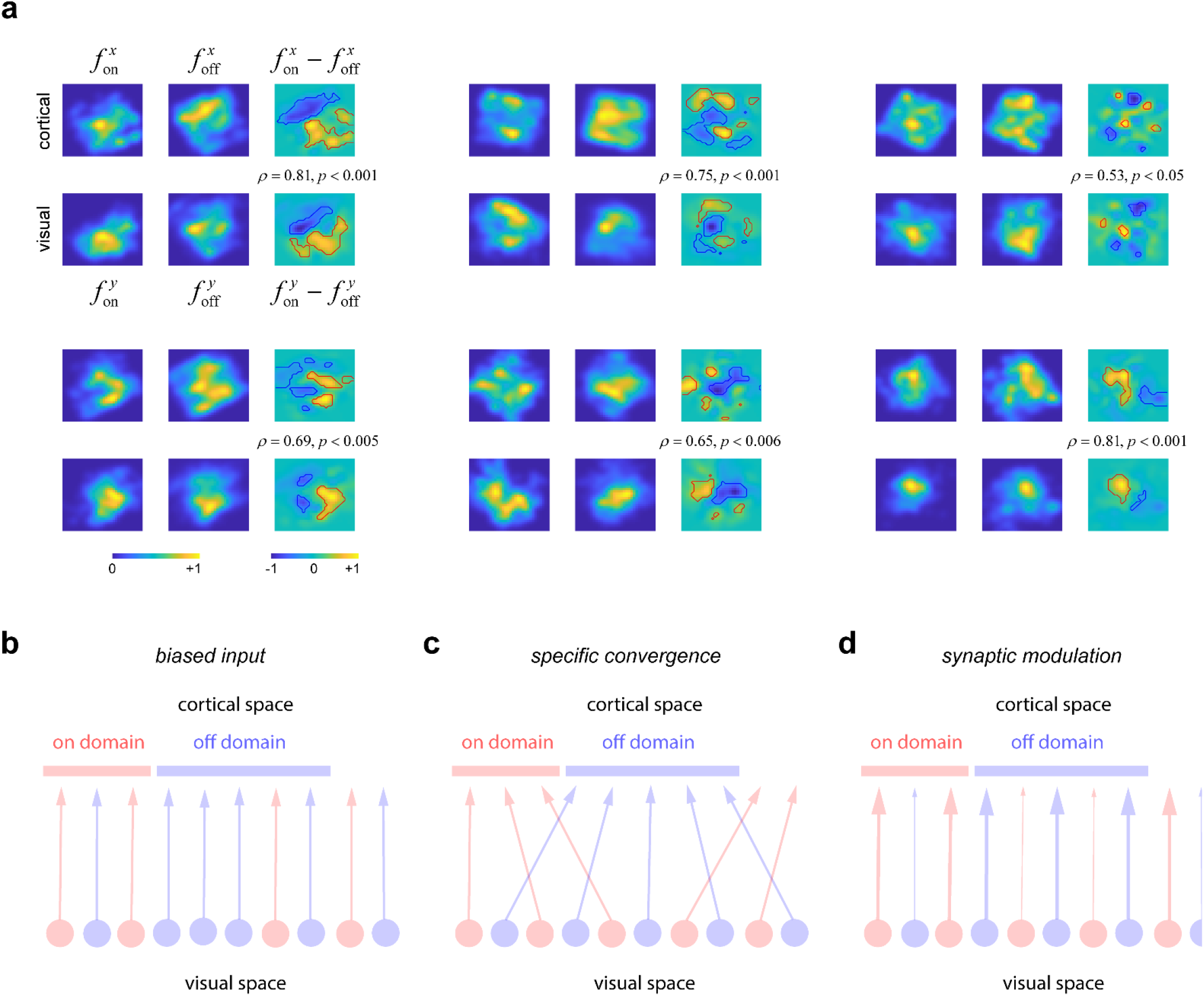
ON/OFF domains in mouse V1. **a,** Each panel displays the result of one experiment. In each case, the top row displays the density of ON 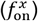 and OFF 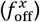 cells in canonical cortical space, along with their difference, 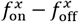. The bottom row shows the density in the position of receptive field centers for ON 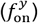 and OFF 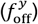 cells in canonical visual space, along with their difference, 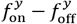 In all panels, both axes span the range from −2.5 to 2.5. Level sets showing areas showing fluctuations above the *p* = 0.01 significance level are shown by red and blue curves. The correlation coefficient between the fluctuations 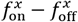 and 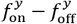 is shown at the inset along with the statistical significance reached. **b-d,** Possible models of ON/OFF domains. **b,** Biased input model. **c,** Specific convergence model. **d,** Synaptic modulation model.

We find that the distributions of 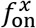 and 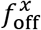 are patchy, peaking in different locations, which results spatial structure in the fluctuations 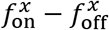 (**Fig. 2a**, top rows in each panel). In other words, some cortical patches contain a higher density of ON cells than OFF cells (ON domains), while others contain higher density of OFF cells than ON cells (OFF domains). We assessed the likelihood that the observed magnitudes in the fluctuations of 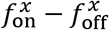 could arise by chance in Monte Carlo simulations where ON/OFF labels were randomly shuffled (**Methods**). Level sets were computed corresponding to the *ρ* = 0.01 significance level (**Fig. 2a**, red and blue solid curves). In all our experiments, we detected significant ON and OFF domains as fluctuations (either positive or negative) exceeding this significance level. We conclude that mouse V1 contains ON/OFF domains.

### Origin of ON/OFF domains

We found that fluctuations in 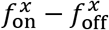 robustly mirrored fluctuations in the density of receptive field centers on the visual field, 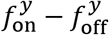 (**Fig. 2a**, bottom row in all panels), as demonstrated by the high correlation between these functions (**Fig 2a**, inset). Statistical significance was established by computing the likelihood that the observed correlations values could arise by randomly shuffling ON/OFF labels (**Fig. 2a**, *p-values*, **Methods**). Note it would have been difficult to notice this relationship between fluctuations in cortical and visual spaces without the representation derived from canonical correlation analysis.

This result constraints models of ON/OFF domain formation. The simplest explanation for the data is that the relative density of ON/OFF receptive field centers at the input varies across the visual field. If the retinotopic mapping is precise enough, such fluctuations would be mirrored in the cortex, generating corresponding ON/OFF domains (**Fig. 2b**, biased input model). Alternatively, it is possible the density of ON/OFF receptive fields is uniform in visual space and that either ON/OFF cells specifically target different domains in the cortex (**Fig. 2c**, specific convergence model) or that the synaptic drive is modulated in space in a way that creates ON/OFF domains (**Fig. 2d**, synaptic modulation model). The specific convergence model would not yield the observed correlations in relative density fluctuation and can be rejected. The synaptic modulation model could explain the data if weak ON inputs into OFF domains (or weak OFF inputs into ON domains) were to be undetectable by our methods. This would result in the appearance that the density of the ON/OFF centers is non-uniform. This scenario predicts that the magnitude of responses of ON cells near the center of OFF domains ought to be lower than the responses of ON cells near the center of ON domains, with a similar prediction applying to OFF cells. However, a two-way ANOVA analysis of responsiveness as a function of cell type (ON/OFF) and domain type (ON/OFF) fails to show any such interaction (*p* > 0.2 in all cases). We can then reject the synaptic modulation model, leaving the biased input model as the most plausible explanation at present.

### ON/OFF domains shape receptive field structure

Next, we asked if there is any relationship between the spatial distribution of ON/OFF domains and the structure of simple cell receptive fields, which integrate both ON and OFF inputs. We denote the average receptive field of ON cells in the entire population by *μ*_on_ and adopt a corresponding definition for *μ*_off_. We define the average of all simple cell receptive fields in the population by *μ*_s_. There is a strong correlation between *μ*_on_ − *μ*_off_ and *μ*_s_ (*p* < 10^−10^ in all cases) (**Fig. 3a**). The average of all the simple cell receptive fields in the population is biased in the same way as the distribution of ON and OFF receptive fields, consistent with the notion that ON/OFF domains shape the structure of simple cells in V1. This result replicates a similar finding in the cat^7^.

**Figure 3.**
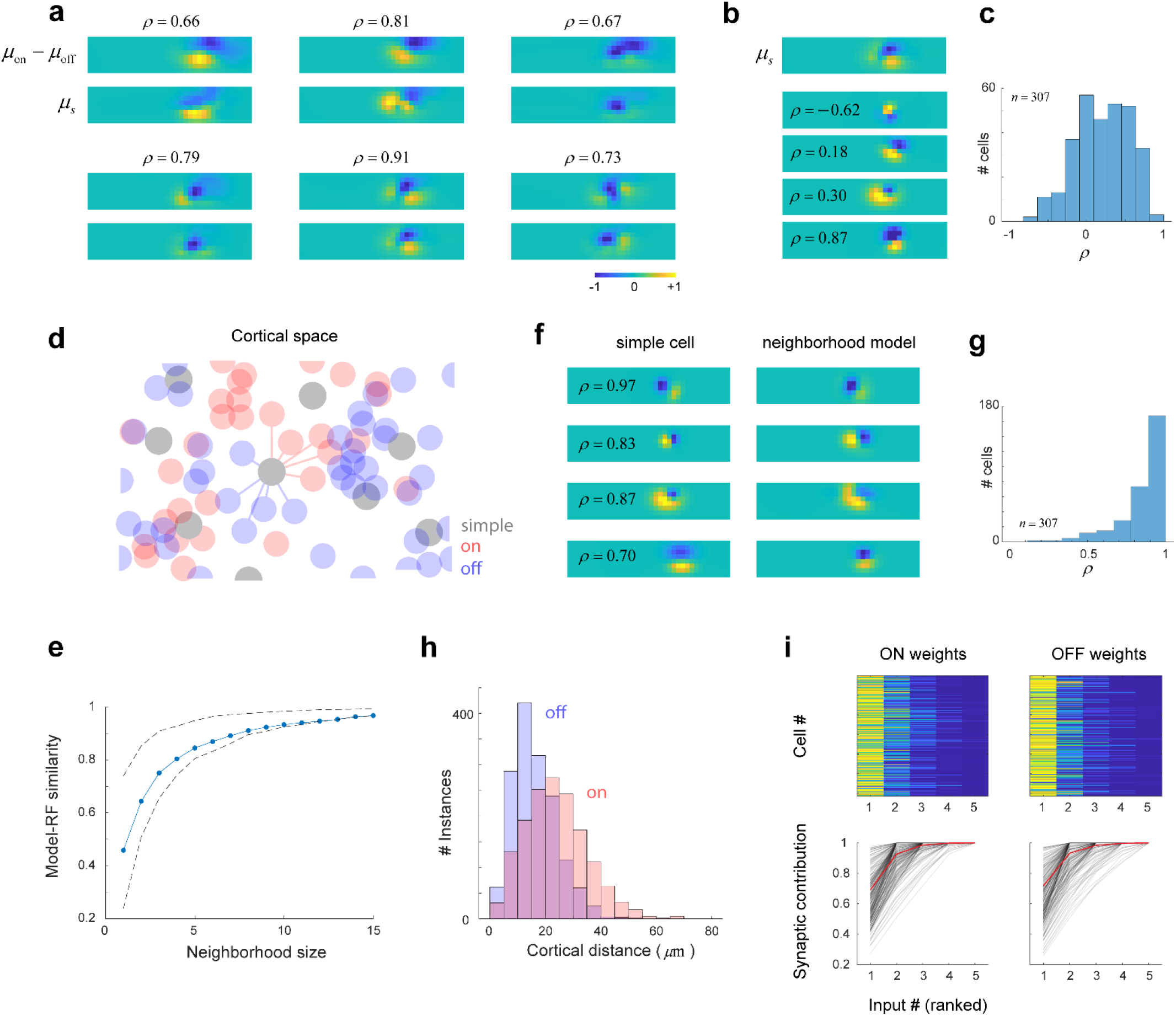
ON/OFF domains shape receptive field structure. **a,** Correlation between ON/OFF and simple cell receptive fields. Each panel displays the result of one experiment. The top row displays the difference between the average ON (*μ*_on_) and OFF (*μ*_off_) receptive fields in the population, while the bottom row shows the average of all simple cell receptive fields (*μ_s_*). The two are correlated (*ρ* < 0.001 in all cases). **b,** Diversity of receptive fields in the population, showing the average along with the receptive fields of individual cells and their correlation with the average. **c,** Distribution of correlation coefficients between *μ_s_* and those of individual cells in one experiment. **d,** Model of simple cells as linear combination of ON and OFF signals in a local neighborhood. 3, Model performance as a function of neighborhood size. Performance starts to saturate at a neighborhood size of *k*~5. Dashed lines indicate 25-th and 75-th percentiles of the distribution of correlation coefficients for each neighborhood size). **f,** Example of model fits. **g,** Model performance for all neurons in one experiment for *k* = 5. **h,** Distribution of distances for ON and OFF from simple cells for a neighborhood size of five – most cells are within 50*μ*m of the target neuron. i, Distribution of weights for ON and OFF neurons in decreasing order by rank (top). Two ON or OFF cells are sufficient to account for 90% of the total synaptic input to the neuron, as shown by the average cumulative distribution (bottom, red curve; individual cells shown by black curves).

Despite the agreement between the averages *μ*_s_ and *μ*_on_ − *μ*_off_, there is substantial variability in the structure of simple cell receptive fields (**Fig. 3b**). This can be observed by computing the distribution of the correlation coefficient between the receptive fields of individual neurons and the average *μ*_s_ (**Fig. 3c**). The variability in the population is reflected in the spread of the correlation coefficient which spans a range from −0.72 to 0.88. The mean of the distribution, of course, is significantly larger than zero (sign-rank test, *p* < 10^−10^) as anticipated from the high correlation between the averages (**Fig. 3a**). Such a pattern of results was typical of all our datasets.

Can the variability in the local distribution of ON and OFF cells account for the diversity of receptive field structures seen in simple cells? To test this idea, we modeled the receptive field of simple cells as a non-negative, linear combination of the receptive fields of its *k*-nearest ON and OFF cells (**Fig. 3d**). To select the size of the neighborhood (the parameter *k*), we calculated how the goodness-of-fit of the model as a function of *k* and found that it begins to saturate at *k*~5 (**Fig. 3e**). The performance of the model for a choice of *k* = 5 was very good: the correlation coefficient between the fits and the actual receptive fields was highly skewed towards one with an average of 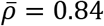 (**Fig. 3f,g**). Cells in the local neighborhood were no more than ~50*μ*m away of the target cell (**Fig. 3h**). Interestingly, there was a large disparity in the weights estimated to the cells in the neighborhood (**Fig. 3i**). When the weights of ON and OFF cells are plotted in descending order, it is evident that their distribution is sparse. On average, more than 90% of the total synaptic input could be accounted for the largest two inputs for both ON and OFF neurons (**Fig. 3i**, bottom, solid dark lines). Such results were typical of our datasets. We conclude that the receptive field of simple cells can be explained by the non-negative, sparse sampling of ON and OFF signals within a neighborhood of ~50*μ*m radius.

## Conclusion

We introduced a novel analysis of the distribution of ON/OFF neurons and the location of their receptive field centers. We showed that, like other mammals^1–4^, mouse V1 is parceled into ON/OFF domains. Our analyses further revealed that fluctuations in the density of ON/OFF neurons on the cortical surface, which defines ON/OFF domains, are mirrored by fluctuations in the density of ON/OFF receptive field centers on the visual field. This finding allowed us to rule out some models of domain formation, leaving the biased input model as the most likely explanation at present.

We further showed that ON/OFF domains shape the receptive field structure of simple cells. First, in each cortical volume, the average of simple-cell receptive fields correlates with the difference between the average of ON and OFF receptive fields. Second, we described how the local diversity of simple-cell receptive fields is explained by a model were neurons linearly combine a sparse number of ON and OFF signals within their cortical neighborhoods^15,20^. Altogether, our findings suggest that ON/OFF domains originate in fluctuations of the spatial density of ON/OFF inputs in the visual field and that they shape the structure of simple receptive fields^10–13,21–23^.

The results lend additional support to the proposal that ON/OFF domains are a critical feature of thalamocortical connectivity influencing the architecture of the cortex^5,14^, and are consistent with the idea that receptive fields in the cortex are constrained by the spatial distribution of ON and OFF inputs in the visual field^10,13,22,27^. While the presence of ON/OFF domains may be sufficient to seed the development of orientation tuning and two-dimensional, simple cell structure, it is clearly not sufficient to establish robust orientation preference maps with the precision observed in higher mammals^17,28^. Instead, orientation columns are maps may additionally require a lower density of thalamic afferents, increased separation between ON and OFF domains, and lower retinotopic scatter of the thalamocortical projection than observed in mice^5,29^.

## Methods

### Experimental model and subject details

All procedures were approved by UCLA’s Office of Animal Research Oversight (the Institutional Animal Care and Use Committee) and were in accord with guidelines set by the U.S. National Institutes of Health. A total of 6 mice, male (3) and female (3), aged P35-56, were used. All these animals were from a TRE-GCaMP6s line G6s2 (Jackson Lab), where GCaMP6s is regulated by the tetracycline-responsive regulatory element (tetO). Mice were housed in groups of 2-3, in reversed light cycle.

### Surgery

Carprofen was administered pre-operatively. Mice were anesthetized with isoflurane (4%–5% induction; 1.5%–2% surgery). Core body temperature was maintained at 37.5C using a feedback heating system. Eyes were coated with a thin layer of ophthalmic ointment to prevent desiccation. Anesthetized mice were mounted in a stereotaxic apparatus. Blunt ear bars were placed in the external auditory meatus to immobilize the head. A portion of the scalp overlying the two hemispheres of the cortex (approximately 8mm by 6mm) was then removed to expose the underlying skull. The skull was dried and covered by a thin layer of Vetbond. After the Vetbond dries (15 min) it provides a stable and solid surface to affix an aluminum bracket (a head holder) with dental acrylic. The bracket is then affixed to the skull and the margins sealed with Vetbond and dental acrylic to prevent infections. Carprofen was also administered post-operatively for 72 hours.

### Two-photon imaging

We conducted imaging sessions starting 6-8 days after surgery. We used a resonant, two-photon microscope (Neurolabware, Los Angeles, CA) controlled by Scanbox acquisition software and electronics (Scanbox, Los Angeles, CA). The light source was a Coherent Chameleon Ultra II laser (Coherent Inc, Santa Clara, CA) running at 920nm. The objective was an x16 water immersion lens (Nikon, 0.8NA, 3mm working distance). The microscope frame rate was 15.6Hz (512 lines with a resonant mirror at 8kHz). The field of view was 900*μ*m × 540*μ*m in all instances. The objective was tilted to be approximately normal the cortical surface. An electronically tuned lens (Optotune EL-10-30-C, Dietikon, Switzerland) was used to run independent sessions acquiring data from optical planes spaced 30*μ*m apart starting at a depth of ~150*μ*m from the cortical surface. A total of 12 datasets were recorded. Images were processed using a standard pipeline consisting of image stabilization, cell segmentation and deconvolution using Suite2p (https://suite2p.readthedocs.io/). For any one optical section, the location of the cells in the imaging plane were estimated as the center of mass of the corresponding region of interest calculated by Suite2p.

### Visual stimulation

We used a Samsung CHG90 monitor positioned 30 cm in front of the animal. The screen was calibrated using a Spectrascan PR-655 spectro-radiometer (Jadak, Syracuse, NY), generating gamma corrections for the red, green and blue components via a GeForce RTX 2080 Ti graphics card. Visual stimuli were generated by a Processing sketch using OpenGL shaders (see http://processing.org). The screen was divided into an 18 by 8 grid (the average size of each tile was 8 by 8 deg of visual angle). Each frame of the stimulus was generated by selecting the luminance of each tile randomly as either bright (10% chance), dark (10% chance) or grey (80% chance). The stimulus was flashed for 166 ms and appeared at a rate of 1 per second. Between stimuli the screen was uniform gray. Transistor-transistor logic (TTL) pulses generated by the stimulus computer signaled the onset of stimulation. These pulses were sampled by the microscope and time-stamped with the microscope frame and line number that was being scanned at that moment. Sessions lasted for 25 min, generating the response of cells in the population to 1500 stimulus presentations.

### Calculation of ON and OFF maps

For each cell and tile in the stimulus we calculated the average response the neuron locked to the presentation of a bright or dark stimulus over the 15 frames (1 sec) following stimulus onset. The ON map of the *k* − *th* cell in the population at a delay of *t* frames after stimulus onset was represented as an image of equal size to the stimulus. The value a tile at location (*i*, *j*)corresponds to the average response of the neuron to the presentation of a bright stimulus at that location *t* frames after stimulus onset. We denote this image by 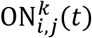 and adopt a similar definition of the OFF map, 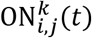. At each we compute the relative norm of the responses (seen as a vector) normalized by the norm at *t* = 0: 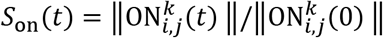 and For each cell and tile in the stimulus we calculated the average response the neuron locked to the presentation of a bright or dark stimulus over the 15 frames (1 sec) following stimulus onset. The ON map of the *k* − *th* cell in the population at a delay of *t* frames after stimulus onset was represented as an image of equal size to the stimulus. The value a tile at location (*i*, *j*)corresponds to the average response of the neuron to the presentation of a bright stimulus at that location *t* frames after stimulus onset. We denote this image by 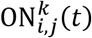 and adopt a similar definition of the OFF map, 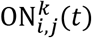. At each we compute the relative norm of the responses (seen as a vector) normalized by the norm at *t* = 0: 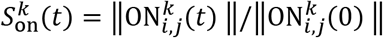 and 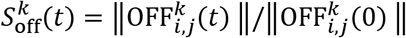. These curves typically peaked at delays of 5 to 7 frames (corresponding to 322-350ms). We declared a cell to have a significant ON map if its normalized norm attained a peak value larger than 5 and a two-dimensional Gaussian fit to the ON map at the peak time had a correlation coefficient to the raw image which was larger than 0.5. A similar definition applied for OFF maps. Cells could have no significant maps, either significant ON or OFF maps, or both. A cell was selected to be part of our analyses if it had at least one significant map. The center of the receptive field was estimated as the center of the Gaussian fit.

### Canonical correlation analysis

Each neuron had assigned a coordinate in cortical space, (*x*_1_, *x*_2_, *x*_3_) (**Fig 1b**) and, for each of its significant maps, one in visual space, (*y*_1_, *y*_2_) (**Fig. 1d**, right panels). Canonical correlation analysis finds transformations 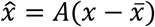 and 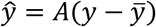 such that the covariance of 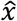 and 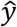 is diagonal and the correlations between matching canonical coordinates are maximized. The transformations are further constrained so that the variance of the canonical coordinates equals one. Note that in our case the matrix *A* is *n* × 3, while the matrix *B* is *n* × 2, where *n* is the total number of cells with at least one significant map.

### Density estimation

Given a distribution of points in canonical coordinate space (either cortical or visual) we estimate the density distribution by 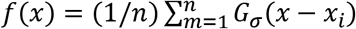, where *G_σ_*(·) is a two-dimensional Gaussian kernel of width *σ* and {*x*_*i*_} (*i* = 1, …, *N*) is the set of points under consideration^30^. We chose a width of *σ* = 0.25, following the rule of thumb bandwidth estimator 0.9 *n*^−1/5^, with *n*~500, which is a typical size for our data^30^. Estimates of 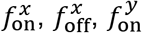 and 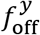 were all obtained by this procedure. Correlation between maps and their statistical significance were calculated using MATLAB (Mathworks, Natick, MA).

To evaluate the likelihood that the observed fluctuations could arise by chance, we randomly shuffled the labels of ON and OFF cells in *N* = 1000 experiments. For each experiment, we calculated the distribution of fluctuations at each point in cortical and visual spaces, which enabled us to compute *p* = 0.01 level sets (**Fig 2**). Similarly, we computed the distribution of correlation coefficients between fluctuations of ON/OFF cells on the cortical surface and those of their center locations in the visual field, allowing us to calculate the statistical significance of the observed correlation in the original data (**Fig 2**).

## Data availability

The data from all experimental sessions are available at____________.

## References

1 Smith, G. B., Whitney, D. E. & Fitzpatrick, D. Modular Representation of Luminance Polarity in the Superficial Layers of Primary Visual Cortex. Neuron 88, 805–818, doi:10.1016/j.neuron.2015.10.019 (2015).

2 Jin, J. Z. et al. On and off domains of geniculate afferents in cat primary visual cortex. Nature neuroscience 11, 88–94 (2008).

3 McConnell, S. K. & LeVay, S. Segregation of on- and off-center afferents in mink visual cortex. Proc Natl Acad Sci U S A 81, 1590–1593, doi:10.1073/pnas.81.5.1590 (1984).

4 Zahs, K. R. & Stryker, M. P. Segregation of ON and OFF afferents to ferret visual cortex. J Neurophysiol 59, 1410–1429, doi:10.1152/jn.1988.59.5.1410 (1988).

5 Mazade, R. & Alonso, J. M. Thalamocortical processing in vision. Vis Neurosci 34, E007, doi:10.1017/S0952523817000049 (2017).

6 Kremkow, J., Jin, J., Wang, Y. & Alonso, J. M. Principles underlying sensory map topography in primary visual cortex. Nature 533, 52–57, doi:10.1038/nature17936 (2016).

7 Jin, J., Wang, Y., Swadlow, H. A. & Alonso, J. M. Population receptive fields of ON and OFF thalamic inputs to an orientation column in visual cortex. Nature neuroscience 14, 232–238 (2011).

8 Lee, K. S., Huang, X. & Fitzpatrick, D. Topology of ON and OFF inputs in visual cortex enables an invariant columnar architecture. Nature 533, 90–94, doi:10.1038/nature17941 (2016).

9 Wang, Y. et al. Columnar organization of spatial phase in visual cortex. Nat Neurosci 18, 97–103, doi:10.1038/nn.3878 (2015).

10 Ringach, D. L. Haphazard wiring of simple receptive fields and orientation columns in visual cortex. Journal of neurophysiology 92, 468–476 (2004).

11 Ringach, D. L. On the origin of the functional architecture of the cortex. PLoS One 2, e251, doi:10.1371/journal.pone.0000251 (2007).

12 Paik, S. B. & Ringach, D. L. Link between orientation and retinotopic maps in primary visual cortex. Proc Natl Acad Sci U S A 109, 7091–7096, doi:10.1073/pnas.1118926109 (2012).

13 Paik, S. B. & Ringach, D. L. Retinal origin of orientation maps in visual cortex. Nat Neurosci 14, 919–925, doi:10.1038/nn.2824 (2011).

14 Kremkow, J. & Alonso, J. M. Thalamocortical Circuits and Functional Architecture. Annu Rev Vis Sci 4, 263–285, doi:10.1146/annurev-vision-091517-034122 (2018).

15 Smith, S. L. & Häusser, M. Parallel processing of visual space by neighboring neurons in mouse visual cortex. Nature neuroscience 13, 1144–1149 (2010).

16 Ohki, K., Chung, S., Ch’ng, Y. H., Kara, P. & Reid, R. C. Functional imaging with cellular resolution reveals precise micro-architecture in visual cortex. Nature 433, 597–603, doi:10.1038/nature03274 (2005).

17 Ringach, D. L. et al. Spatial clustering of tuning in mouse primary visual cortex. Nat Commun 7, 12270, doi:10.1038/ncomms12270 (2016).

18 Ji, W. et al. Modularity in the Organization of Mouse Primary Visual Cortex. Neuron 87, 632–643, doi:10.1016/j.neuron.2015.07.004 (2015).

19 Niell, C. M. & Stryker, M. P. Highly selective receptive fields in mouse visual cortex. Journal of Neuroscience 28, 7520–7536, doi:10.1523/jneurosci.0623-08.2008 (2008).

20 Ringach, D. L. Sparse thalamocortical convergence. Current Biology 31, 2199–2202. e2192 (2021).

21 Jimenez, L. O., Tring, E., Trachtenberg, J. T. & Ringach, D. L. Local tuning biases in mouse primary visual cortex. J Neurophysiol, doi:10.1152/jn.00150.2018 (2018).

22 Ringach, D. L. in The New Visual Neurosciences Vol. The New Visual Neurosciences (eds J. S. Werner & L. M. Chalupa) (2013).

23 Ringach, D. L. You get what you get and you don’t get upset. nature neuroscience 14, 123–124 (2011).

24 Hubel, D. H. & Wiesel, T. N. Receptive fields, binocular interaction and functional architecture in the cat’s visual cortex. J Physiol 160, 106–154 (1962).

25 Skottun, B. C. et al. Classifying simple and complex cells on the basis of response modulation. Vision Res 31, 1079–1086 (1991).

26 Hotelling, H. Relations between two sets of variates. Biometrika 28, 321–377 (1936).

27 Soodak, R. E. The retinal ganglion cell mosaic defines orientation columns in striate cortex. Proc Natl Acad Sci U S A 84, 3936–3940 (1987).

28 Bonin, V., Histed, M. H., Yurgenson, S. & Reid, R. C. Local diversity and fine-scale organization of receptive fields in mouse visual cortex. J Neurosci 31, 18506–18521, doi:10.1523/jneurosci.2974-11.2011 (2011).

29 Jang, J., Song, M. & Paik, S.-B. Retino-cortical mapping ratio predicts columnar and salt-and-pepper organization in mammalian visual cortex. Cell reports 30, 3270–3279. e3273 (2020).

30 Silverman, B. W. Density Estimation for Statitics and Data Analysis. (Chapman & Hall, 1986).

